# Rare plants can make an important contribution to sustain local biodiversity through biological interactions

**DOI:** 10.64898/2026.05.16.725624

**Authors:** María B García, Héctor Miranda-Cebrián, Miguel Verdú, David Martín, Javier Blasco-Zumeta, María Jarne, Jens Olesen

**Author notes:** Both authors contributed equally (María B García, Héctor Miranda-Cebrián).

## Abstract

Plants, as structural elements of habitats, contribute greatly to the maintenance of local biodiversity through their biological interactions. In this study we explore whether their rarity, according to Rabinowitz’s (1981) three criteria, is related to the richness and diversity of arthropods and other plants they are associated to, in a gypsum-rich steppe. We first analysed whether the geographic abundance and ecological specialisation of 32 characteristic and dominant plant species are related to the diversity (richness and phylogenetic diversity (MPD)) and degree of local specialisation of arthropods associated with them (1,694 taxa). Then, we focused on a non endemic and non specialized plant in the study area (*Krascheninnikovia ceratoides*) to explore the effect of population size on two types of interactions: aerial arthropods and plant facilitation. Results indicate that: 1) plant species abundance (geographical range) is not related to the richness or MPD of communities of associated arthropods, 2) plant species ecological specialization (edaphic endemisms or ‘gypsophiles’) do not contribute differentially to the maintenance of singular arthropod communities, and 3) the community of aerial arthropods and plants interacting with *K. ceratoides* in a small population are not necessarily less diverse than those in patches of similar size in a large population. Results also revealed that the two plant species with fewer interactions (one rare, one widespread) do show the highest singularity in their interactions with arthropods. Our study illustrates the important contribution of rare plants to the conservation of local biodiversity.

## INTRODUCTION

Biological interactions are key to biodiversity. Not only have they played an important role in its past generation, but living organisms today depend on them for their persistence (Bascompte and Jordano 2007). Although interactions are often overlooked and not easy to record in the field because of their short duration (e.g., pollination) or the small size of the interacting organisms (e.g., mycorrhizal fungi), they are the necessary “glue” that keeps species and communities together and functioning in natural systems. Therefore, the more interactions a species is involved in, the more biodiversity it helps to maintain.

Interactions can determine the abundance or set the range limits of species (Louthan et al. 2015), and their significance can be such that the loss of a single interaction can result in the malfunction of any of the organisms involved, and even precede its local extinction (Valiente-Banuet et al. 2015). Similarly, the disappearance of a single species from an ecosystem can have important consequences for the maintenance of overall biodiversity through the interactions in which it is involved, either directly or through indirect cascading effects (Rezende et al. 2007; Dirzo et al. 2014). In this context, it seems important to explore the extent to which the vulnerability of one partner may affect the interactions in which it is involved.

Rarity is often a signal of vulnerability because it places species in situations where chance (e.g. demographic stochasticity due to small population sizes) may play a negative role. In addition, rare species seem to have less scope for recovery and lower plasticity to cope with disturbance (Kempel et al. 2020). Consequently, interactions involving rare species may have a higher risk of disappearance, as the local persistence of a rare species may determine the fate of other associated rare organisms through complex multitrophic interactions (Kéry et al. 2001). Rarity, however, is a complex and multidimensional concept. Four decades ago, Rabinowitz (1981) proposed a system for classifying species rarity that is still relevant today, based on three axes: small geographic range, ecological specialisation and small population size. Reduced distribution (typical of regional or local endemics) or territorial abundance usually means that the species is very sensitive to factors that can lead to its disappearance by chance or by negative factors acting at a regional or local scale (Purvis et al. 2000b; Payne and Finnegan 2007; Humphreys et al. 2019). Ecological specialisation can become another serious negative factor as it can lead to reduced competitive ability and restrict the species’ occurrence to discrete and infrequent patches (Walker and Preston 2006; Zettlemoyer et al. 2019). Finally, low numbers of individuals in populations may bring to the fore the importance of stochastic processes such as the founder effect or the Allee effect (Pimm et al. 1988; Matthies et al. 2004; O’Grady et al. 2004). All these factors have been identified as drivers of extinction in several large-scale studies (Purvis et al. 2000a; Harnik et al. 2002; Davies et al. 2004; Saupe et al. 2015). While rare species have been the focus of numerous conservation studies in the past (e.g., Bakker and Doak 2009; Gerber 2016; Enquist et al. 2019), and the study of biotic interactions has developed considerably over the last two decades (e.g., Bascompte et al. 2003; Landi et al. 2018; Karban and Agrawal 2023), studies combining rare species and biotic interactions are scarce (but see Kempel et al. 2020).

Plants are essential nodes of ecological networks, interacting with co-occurring plants and a wide variety of animals, from pollinators to criptic below ground organisms such as detritivores or consumers through predation on seeds (Braun et al. 2021), which influence ecosystem functioning. An interesting question, therefore, is to explore the overall contribution of rare plants to maintain arthropod diversity compared to co-occurring more common or generalist plant species in the same location. Using a phytocentric and integrative approach (see Olesen 2022), in this study we focus on the effect of rarity of plant species on interactions along the three proposed axes (geographical extent, ecological specialisation, and population size), and therefore on the local biodiversity they help to maintain.

Our study area is a semi-desert where plants experience high levels of calcium and sulphate, and low availability of water and organic matter, an stressful environment that has favoured the occurrence of specialised plants. These “gypsophiles” are able to cope with the physical and chemical limitations imposed by gypsum soils, coexisting with more generalist plants (gypsovages; Palacio et al. 2007; Escudero et al. 2015). This context gave us the opportunity to test the effect of rarity on biological interactions resulting from ecological specialization in addition to the effect of geographical extent. We first explored the taxonomic richness and evolutionary diversity of above- and below-ground arthropod community associated with 32 plant species typically characteristic of this type of habitat, and also for the local specialization of arthropods to each plant species. Then, for the effect of population size, we focused on a non-endemic and non-specialist plant species of the community, *Krascheninnikovia ceratoides*, and two types of interactions (plant facilitation, and interactions with aerial arthropods) in one small population and three patches of similar size in a large population.

Our goal is to investigate whether the rarity of plant species in a southern European Mediterranean gypsum-rich steppe is correlated with the richness, diversity and local specialisation of arthropod and plant communities with whom they are associated. Our aim is to provide a general insight into the effect of host plant abundance on maintaining local diversity, in order to reveal potential rarity-driven asymmetries. Investigating whether rare plants show any pattern in their biological interactions is important in the current context of global change, particularly in rare or vulnerable habitats, as the degradation and fragmentation of habitats and the concomitant decline in populations often have negative, but still unpredictable, consequences for those involved (Tylianakis et al. 2008).

## METHODS

### Study area and plant species sampled

The study was carried out in a steppe zone of the Ebro valley, NE of the Iberian Peninsula (Fig. 1). The natural habitat is embedded in a mosaic of agricultural landscapes, with patches of steppe of different sizes remaining in hilly areas. The ecological value of the study area has lead to its inclusion in the Natura 2000 network (LIC/ZEPA Monegros). Nevertheless, the area is currently used for hunting, agriculture and livestock farming. As a result, the natural vegetation has been displaced to areas difficult to clear.

**Fig. 1.**
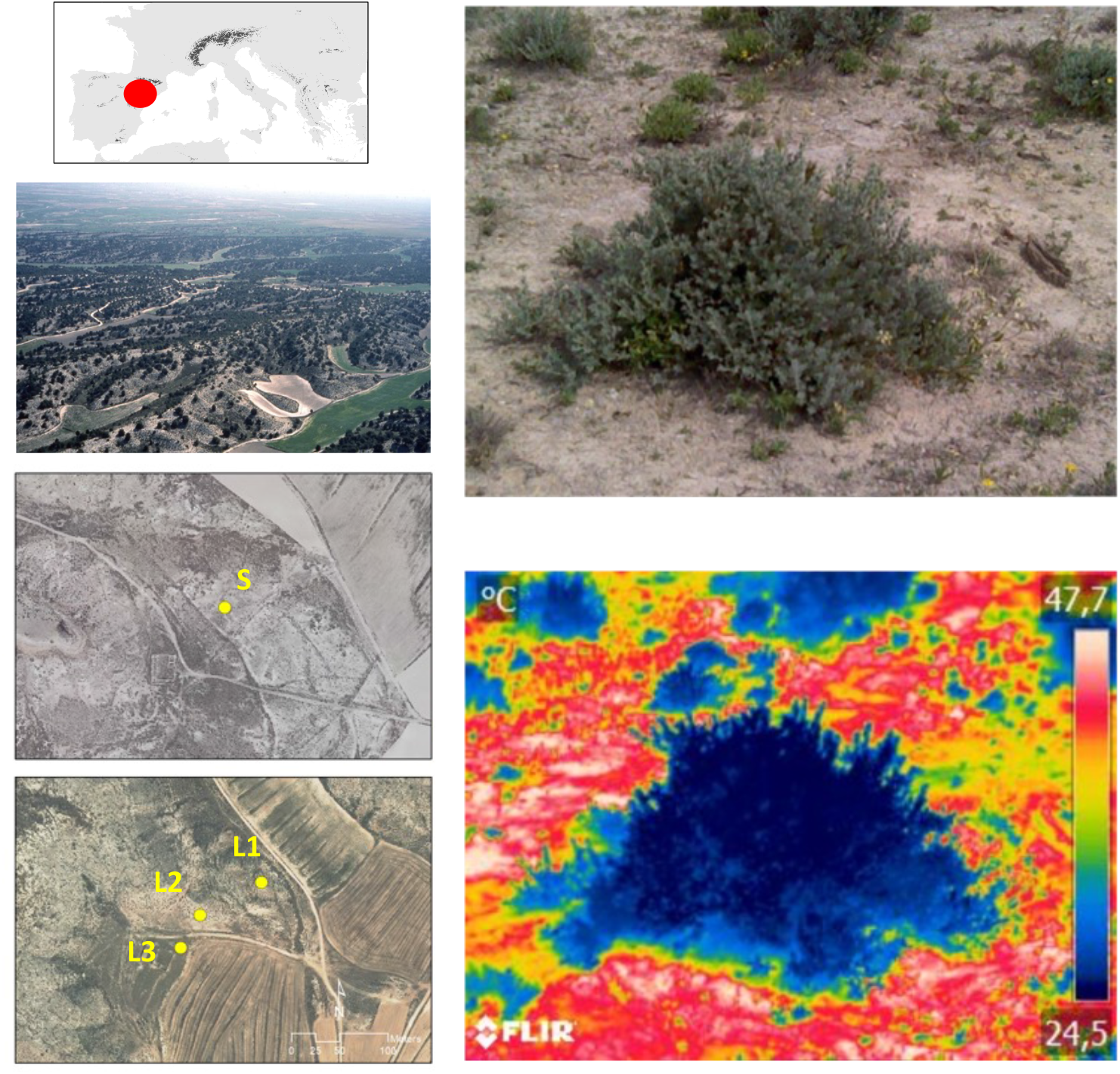
Left: General location of the study area in western Europe (above), aerial picture showing the study area (middle), and populations of Krascheninnikovia ceratoides (below): small (S) and 3 patches of the large population (L: L1, L2 and L3). Right: RGB (above) and thermal (below) images of an individual taken on a day of high insolation (midday, May 2018). Note the temperature contrast between the interior of the plant, the shaded area at the outer edge, and the bare ground around it.

The climate is dry, semi-arid and cold according to the Köppen climate classification. Hence, the cold winters (mean yearly minimum temperature ∼3 ºC) and dry, hot summers (mean yearly maximum temperature ∼ 33ºC and <500 mm per year with high inter-annual variability), together with the high evapotranspiration potential, have created a vegetation with markedly xerophytic character. The soil is very rich in sulphates and calcium, poor in nutrients, sometimes with high salinity and often with surface crusting due to intense evaporation, making it a difficult environment for many plants. Mediterranean plants, both annual and woody, dominate, and specialists exclusive to the gypsum-rich soils coexist with ruderal and arborescent plants. The woody plants often create a microclimate under their shade or among their foliage that facilitates the life of numerous other plants and invertebrates (Fig. 1). The invertebrate fauna of the study area has been intensively studied over the last three decades, with more than 3,500 species been described by more than 100 taxonomists worldwide, including a significant number of regional endemics (Pedrocchi 1998; Blasco-Zumeta 2023).

To test the effect of rarity driven by plant geographic and ecological specialization we chose 32 perennial plants characteristic of a gypsum formation of about 2,000 hectares of juniper (*Juniperus thurifera* L.) located in this area (“La Retuerta de Pina”; 300-400 m a.s.l.; UTM 30TYL29). The visitors they receive were identified and analysed between 1987 and 2000 (Blasco-Zumeta 2023). To test the effect of rarity driven by population size, we conducted a comparative study between 2016 and 2018 of a non-endemic and non-specialist plant of the community, *Krascheninnikovia ceratoides* (L), a relict shrub with an Iranian-Turanian distribution that finds its western limit in the Iberian Peninsula and has affinity for gypsum soils (Fig. 2). This is an anemophilous shrub with dry fruits and hard, hairy leaves usually browsed by sheep. Given the difficulty of finding small populations in the study area, we restricted our sampling to one isolated patch of ∼800 m^2^ and ∼100 individuals (hereafter ‘S’) and a very large population nearby (12 km distance; hereafter ‘L’, containing several thousand plants) to make sure they both experimented very similar climatic and edaphic condictions. Since a larger population will contain more interactants simply because it is spread over a higher environmental heterogeneity than a small one (Galiana et al. 2022; Blasco-Zumeta 2023), to make a proper comparison, we selected and sampled three patches of similar size to the small population inside the large population (L1, L2 and L3; Fig. 1), located at a distance of at least 500 m from each other. Therefore, our study does not make a direct comparison between a large and a small population. Instead, we compare patches within the large population that are similar in size to the small population, ensuring the same sampling effort was applied to the small population and all the three patches of the large one. To confirm that the density of the focal plant in the two study populations was similar too, we applied the point intercept method every 20 cm along several transects (300 contact points along two transects 30 m long in S; 1000 points along 4 transects 50 m long in L), which resulted in very similar density in both populations: 12.3% of contacts in S and 12.5% in L.

**Fig. 2.**
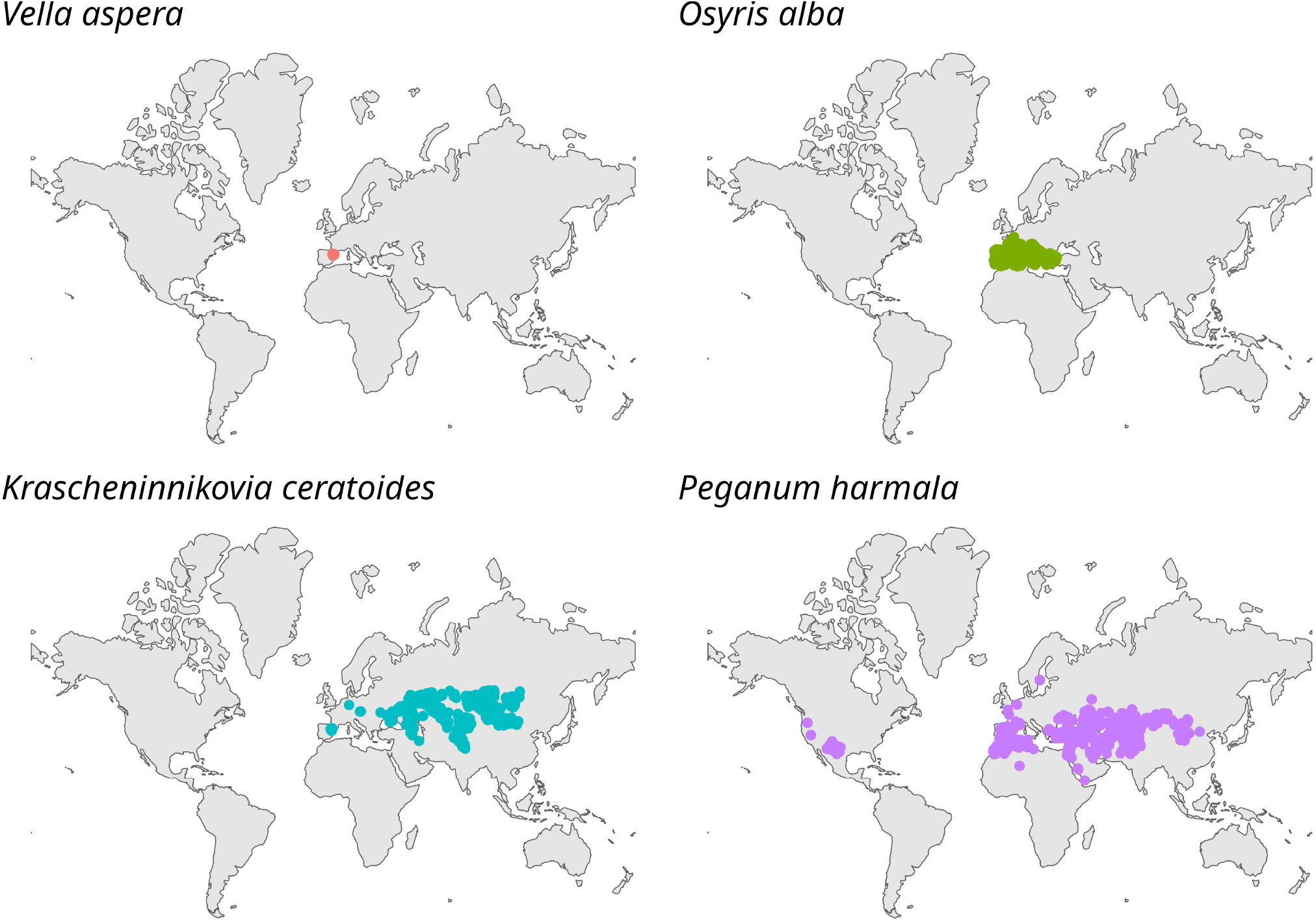
Geographical distribution of four focal steppe plants, showing the wide diversity of occupancies according to GBIF records: from regional endemics to species found on four continents.

### Sampling plants and arthropods associated to focal plants

Different sampling methods were used to describe and quantify the richness and diversity of arthropods associated with the 32 dominant steppe plant species (focal plants thereafter). Details of sampling can be found in Blasco-Zumeta (2023), but basically four types of methods were used: vacuuming with an entomological aspirator and shaking of shoots and branches on plastic bags every 15 days over a year, Wilkening traps inside branches once every 15 days over a year, Berlesse funnels to extract arthropods from the soil below the plant once per season over a year, and emergence boxes to capture insect larvae in fruits, inflorescences and other plant structures. All plants were sampled throughout the year except *Lithospermum fruticosum*, which was sampled only from April to September, and *Peganum harmala*, which was sampled only from March to July. A variable number of (N>10) plant individuals were used for each sampling in order to better cover the study area. Thanks to a network of international collaborators, identification of arthropods to species level was possible in almost all cases. The complete list of species, the higher category (order/subfamily) and the plant species in which they were recorded can be found in Blasco-Zumeta (2023).

For *K. ceratoides* we used two different sampling methods. Aerial arthropod sampling was carried out in spring and autumn by shaking the shoots of a total of 168 plants of different sizes in plastic bags (21 per season in each of the 4 patches: S, L1, L2 and L3) and all specimens were subsequently identified with the help of specialists (see list of taxa in García et al. 2023). Plants facilitated by the microclimatic environment this species creates (growing under their shade or within their foliage) were recorded for 10 *K. ceratoides* individuals of different sizes (3 small, 3 medium, 4 large) distributed along a transect within each patch (S, L1, L2, L3).

### Evolutionary diversity of plants and insects

To characterise the evolutionary relationships (*sensu* Rangel et al. 2015) between the plants of the steppe community and the plants facilitated by *K. ceratoides*, a phylogeny was constructed using the V.PhyloMaker2 package (Yi and Qian, 2022), based on the GBOTB phylogeny. 54 of the 112 species in our study are not present in the phylogeny, so we included them according to taxonomic criteria: those with congeners in the phylogeny (48 out of 54) were grafted next to them, and in the case of only species of the same family but without congeners (6 out of 54), they were grafted randomly within their family. To reduce the phylogenetic uncertainty introduced by this method, 100 different versions of the phylogeny were generated to account for different evolutionary hypotheses *sensu* Rangel et al. 2015).

The evolutionary relationship of the arthropods community was restricted to the class Insecta (88.13% of the recorded arthropods). A similar procedure to the plant phylogeny was followed for the insect phylogeny, in this case using the phylogeny of Chesters (2020). This phylogeny is the most comprehensive published to date for this group of animals, containing more than 100,000 taxa. The evolutionary diversity of the insect communities associated with each focal plant of the steppe was estimated using the mean phylogenetic distance between pairs of species in each community (MPD). Since this metric is highly correlated with the richness of the community under study, making it difficult to compare between different communities we calculated its standard effect size (MPD_SES_) using the ses.mpd function in the picante package (Kembel et al. 2010). This function standardizes the observed MPD value using the mean MPD and its standard deviation obtained by randomizing species across communities but keeping community richness constant. In the case of *K. ceratoides*, this procedure was repeated for the insect and plant communities associated with each of the population patches studied.

### Linking rarity to richness and evolutionary diversity of associated plants and animals

For each steppe focal plant, the global geographic extent of occurrence and area of occupancy (AOO) were calculated from records provided by GBIF (https://www.gbif.org). Only records of georeferenced direct field observations were downloaded. To ensure that these observations were representative of the distribution of each species, we estimated an ellipse containing 99% of these observations and discarded any records that fell outside of it. The geographic range of each species was calculated as the area of the minimum convex polygon (MCP) formed by the selected observations. To avoid possible overestimation of the geographic extent of each species due to overlap of the MCP with marine areas, the surface of the MCP was trimmed with a layer of continents to exclude marine areas. The area of occupancy (AOO) of each focal plant was calculated by counting the number of 2×2 km global grid squares in which each species occurred, based on the observations used to calculate the geographic extent of each species. As both metrics gave similar results, we will refer to the AOO for simplicity throughout the text when testing the geographic abundance effect. Each focal plant was also assigned a value as ‘specialist’ if restricted to gypsum, according to the ‘gypsophily’ value provided by Mota et al. (2011). We also calculated the “arthropod local specialisation” for each focal plant, understood as the average percentage of the 32 focal plants associated to each arthropod. High values correspond to “generalist” arthropods (associated to many plants), and low values to “specialist” arthropods (singular associations). Thus, for each focal plant the final dataset included: area of extent, occupancy (AOO), number of visiting arthropods, evolutionary diversity of such arthropod community, and arthropod local specialization (Table 1).

**Table 1.** Focal plant species in the steppe community of the study area, acronym, family, extent of occurrence, area of occupancy (AOO), gypsophily value for especialised plant (>3.5), number of recorded interactions with aerial arthropods, standardized evolutionary diversity of the arthropod community interacting with each plant species, and mean degree of specialization of such community (see details on how it is estimated in the text). Sampling methods: S: Shake of shoots on plastic bags and entomological hose; B: Berlesse funnel; E: Emergence boxes; W: Wilkening trap inside branches.

| Plant name | Acronym | Family | Extent of occurrence (km<sup>2</sup>) | Occupancy area (number of cells 2x2 km) | Gypsophily value | Sampling methods | N taxa arthropods | MPD.obs.s.z |
| --- | --- | --- | --- | --- | --- | --- | --- | --- |
| *Artemisia herba-alba* Asso | A_her | Asteraceae | 676706 | 709 |  | S, B, E | 294 | 3.782 |
| *Asparagus acutifolius* L. | A_acu | Asparagaceae | 2026095 | 3655 |  | S, E | 29 | 0.805 |
| *Atriplex halimus* L. | A_hal | Amaranthaceae | 8965694 | 1423 |  | S, B, E | 151 | 2.332 |
| *Brachypodium retusum* (Pers.) Beauv. | B_ret | Poaceae | 882349 | 2272 |  | S, B, E | 127 | 3.295 |
| *Carduus bourgeanus* Boiss. & Reuter | C_bou | Asteraceae | 552831 | 419 |  | S | 187 | -2.817 |
| *Caroxylon vermiculatum* (L.) Akhani & Roalson | C_ver | Amaranthaceae | 1840784 | 560 |  | S, B, E | 169 | 2.498 |
| *Centaurea calcitrapa* L. | C_cal | Asteraceae | 166085176 | 3561 |  | S, E | 51 | -0.816 |
| *Ephedra nebrodensis* Tineo ex Guss. | E_neb | Ephedraceae | 488786 | 433 |  | S, B, E | 123 | -0.578 |
| *Frankenia thymifolia* Desf. | F_thy | Frankeniaceae | 183767 | 143 |  | S, B | 53 | 0.824 |
| *Genista scorpius* (L.) DC. | G_sco | Fabaceae | 792618 | 2562 |  | S, E | 71 | 1.505 |
| *Gypsophila struthium* L. ssp. *hispanica* (Willk.) G. López | G_str | Caryophyllaceae | 202379 | 317 | 4.69 - 4.77 | S, B | 286 | -0.710 |
| *Helianthemum squamatum* (L.) Pers. | H_squ | Cistaceae | 231784 | 375 | 4.87 | S, B | 48 | 0.130 |
| *Juniperus phoenicea* L. | J_pho | Cupressaceae | 1630445 | 2203 |  | S, B, E, W | 132 | 2.714 |
| *Juniperus thurifera* L. | J_thu | Cupressaceae | 843191 | 740 |  | S, B, E, W | 427 | 3.397 |
| *Krascheninnikovia ceratoides* (L.) Guedl. | K_cer | Amaranthaceae | 11641072 | 438 |  | S | 9 | 1.083 |
| *Lepidium subulatum* L. | L_sub | Brassicaceae | 296009 | 264 | 4.91 | S | 85 | -1.130 |
| *Lithodora fruticosa* (L.) Griseb. | L_fru | Boraginaceae | 739134 | 1506 |  | S, B | 27 | 1.681 |
| *Lygeum spartum* L. | L_spa | Poaceae | 417888 | 562 |  | S | 10 | 0.632 |
| *Ononis tridentata* L. | O_tri | Fabaceae | 325413 | 623 | 4.13 - 4.53 | S, B | 73 | 3.101 |
| *Osiris alba* L. | O_alb | Santalaceae | 3205568 | 3019 |  | S | 58 | -0.748 |
| *Peganum harmala* L. | P_har | Tetradiclidaceae | 41100598 | 938 |  | S | 29 | 1.733 |
| *Pinus halepensis* Miller | P_hal | Pinaceae | 2865455 | 3457 |  | S, B, E, W | 250 | 0.333 |
| *Quercus coccifera* L. | Q_coc | Fagaceae | 3217835 | 3721 |  | S, B, E | 114 | 1.217 |
| *Retama sphaerocarpa* (L.) Boiss. | R_sph | Fabaceae | 232348 | 1493 |  | S, E | 171 | -3.167 |
| *Rhamnus lycioides* L. | R_lyc | Rhamnaceae | 1342012 | 1778 |  | S, E | 67 | 0.196 |
| *Rosmarinus officinalis* L. | S_lav | Lamiaceae | 47190 | 103 |  | S, B, W | 47 | 0.142 |
| *Salvia lavandulifolia* Vahl. | S_ros | Lamiaceae | 2039570 | 4820 |  | S, B | 241 | -0.195 |
| *Santolina chamaecyparissus* L. ssp. *squarrosa* (DC.) Nyman | S_cha | Asteraceae | 9256505 | 3371 |  | S, B | 70 | 2.723 |
| *Suaeda vera* J.F. Gmelin | S_ver | Amaranthaceae | 5173799 | 800 |  | S, B | 148 | 4.264 |
| *Tamarix canariensis* Willd. | T_can | Tamaricaceae | 678208 | 811 |  | S | 257 | -8.821 |
| *Thymus vulgaris* L. | T_vul | Lamiaceae | 3421150 | 5874 |  | S, B, E | 41 | 0.496 |
| *Vella aspera* Pers. | V_asp | Brassicaceae | 4875 | 29 |  | S | 8 | 1.123 |

Prior to further analyses, the phylogenetic signal of the richness of arthropod-associated community with each focal plant and their geographic abundance was examined using Blomberg’s K to rule out possible effects due to plant relatedness. Since no phylogenetic signal was detected (Mean K = 0.074, SD = 0.004), LMs and GLMs were used to test the relationship between plant rarity (AOO) and arthropod community richness, as well as the relationship between the AOO of each focal plant and the MPD_SES_of insects in their associated communities. We also examined the relationship with the rarity of the interactions they support, or arthropod local specialization.

The relationship between the phylogenetic distance of plants and that of insect communities was analysed using a Mantel test between the average phylogenetic distance matrices between plants (given that there are 100 versions of each phylogeny) and a distance matrix for insect community composition calculated using Sørensen’s index. A non-metric multidimensional scaling (NMDS) analysis was also used to explore the uniqueness of visiting arthropod communities and their relationship with plant ecological specialisation.

In the case of *K. ceratoides*, for which we tested the effect of patch size on richness and MPD_SES_of associated arthropod communities and facilitated plants, possible differences between the small (S) and the large patches (L1, L2 and L3) were estimated by calculating the standard error of the mean of the three large patches, and checking whether the S value was included in the 95% confidence interval of the large population.

## RESULTS

The 32 steppe focal plant species are found along a wide gradient of abundance according to their global distribution and occupancy: from the rarest endemic to the study area (*Vella aspera*) to plants distributed over four continents (*Peganum harmala, Centaurea calcitrapa*; Table 1 and Fig. 2). In terms of ecological specialisation, a small part of this flora (12.5%) consists of plants that only occur in soils with a high concentration of gypsum (gypsophiles; gypsophilic values > 3.5; Table 1).

The sampling of aerial arthropods revealed a total of 1,694 taxa and 3,853 different associations (plant-arthropod combinations). Insects were the main visitors (88.1%), followed by arachnids (8.4%; Fig. 3). Common trees and shrubs such as *Juniperus thurifera, Tamarix canariensis* or *Gypsophila struthium* ranked very high in terms of the number of species associated (≥250 arthropods each), while three focal plants were associated with ≤10 arthropods (Table 1).

**Fig. 3.**
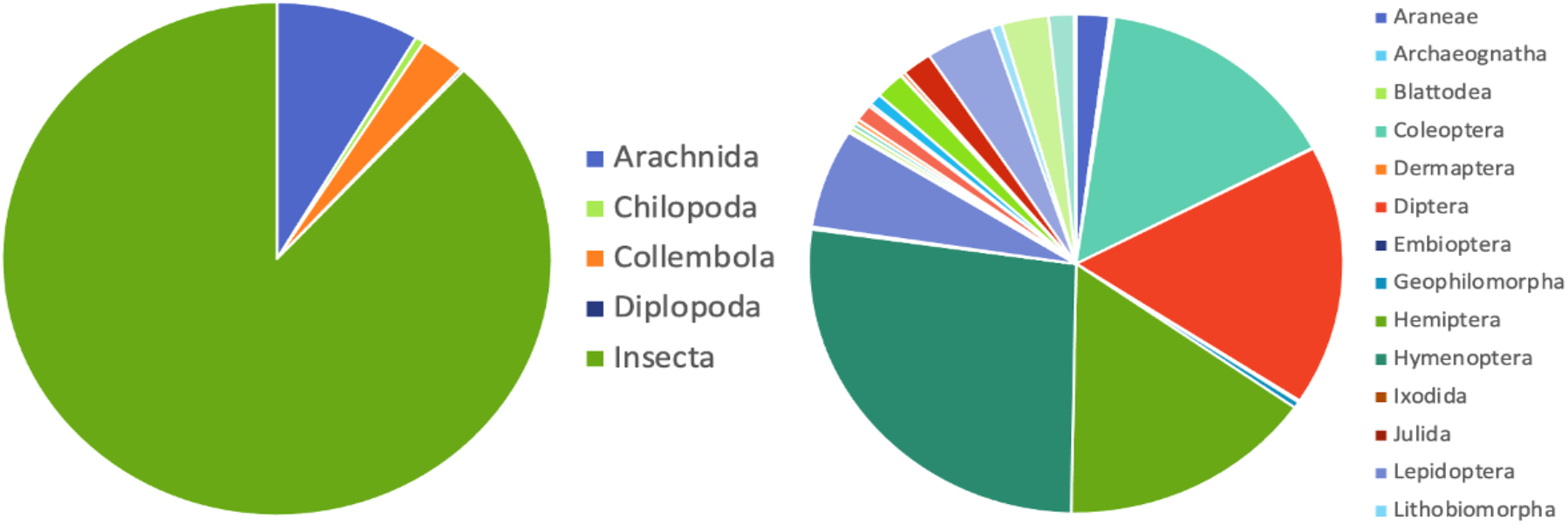
Abundance of different arthropod groups (N=1,697 species) recorded during sampling of focal plants of a gypsum-rich steppe of the Iberian Peninsula.

The area of occupancy (AOO) of steppe plants did not have a statistically significant association with the richness of the arthropods communities they support (GLM with negative binomial response to control for statistical overdispersion; p=0.683). The occupancy area was not associated with the evolutionary diversity of arthropod communities (MPD_SES_; R^2^: -0.032, p=0.845). Therefore, a correlation between plant rarity in geographical terms and the richness or evolutionary diversity of the arthropod communities is ruled out. The NMDS revealed that gypsophiles support arthropod communities relatively similar to those of other ecologically unspecialized plants (Fig. 4).

**Fig. 4.**
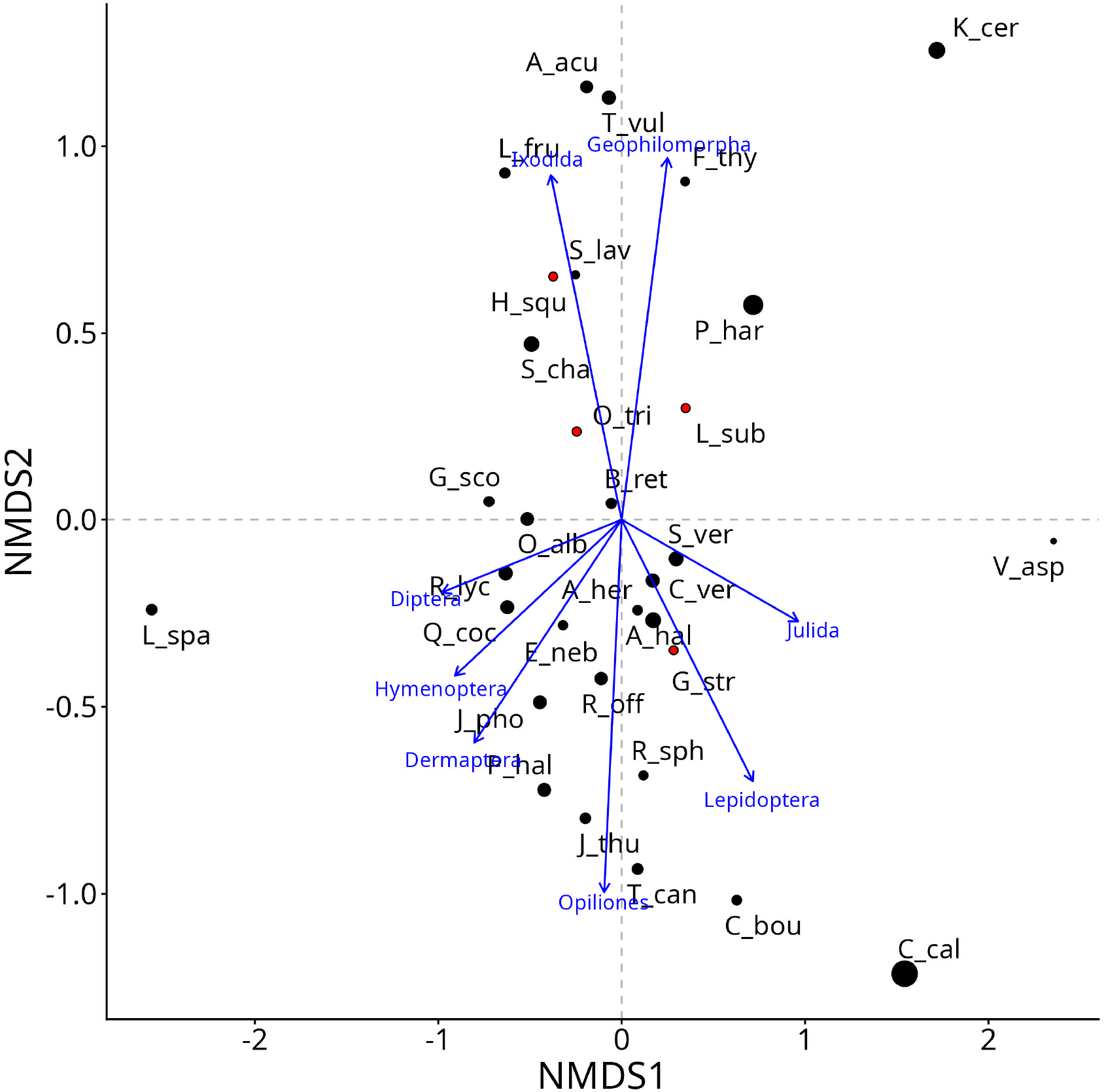
Relative position of each focal plant in the steppe area of study in a Non-metric multidimensional scaling analysis (NMDS), in order to better visualize similarities between the arthropod communities interacting with them. Plant acronyms are listed in Table1. Red dots correspond to gypsophile plants. The size of dots represents total occupancy area (AOO). Blue arrows correspond to the largest taxonomic groups of arthropods.

The relationship between the geographic rarity of plants (AOO) and the rarity of interactions in which they participate (mean % of community plants also visited by their interactants) was also non significant (R^2^ = -0.031, p=0.811). Three plants showcase that lack of correlation: *Vella aspera, Krascheninnikovia ceratoides* and *Lygeum spartum* have interactions with the lowest number of arthropods in the community (8, 9 and 10 arthropods respectively), which are highly specialised on them, while these plants have a variety of geographical extensions (from endemic to pluriregional).

The effect of population size on facilitated plants and aerial arthropods interacting in *K. ceratoides* is shown in Fig. 5. The patch of the small population (S) hosted a richer community of arthropods and they were slightly less phylogenetically related to each other than in the 3 patches of similar size of the large population (L1, L2, L3), although the values in both populations indicate phylogenetic overdispersion (i.e., higher phylogenetic diversity than expected by chance). In the case of facilitated plants, the situation was the opposite: the number of plants facilitated by *K. ceratoides* individuals in the small population was lower (S=25) than in all the three patches inside the large population (L1=39, L2=30, L3=33), and the value of S is outside the confidence interval of the three patches of L. The set of facilitated plants in S is slightly less phylogenetically clustered, although in both cases the negative values of MPD_SES_ indicate a strong tendency towards phylogenetic clustering (i.e., lower phylogenetic diversity than expected by chance).

**Fig. 5.**
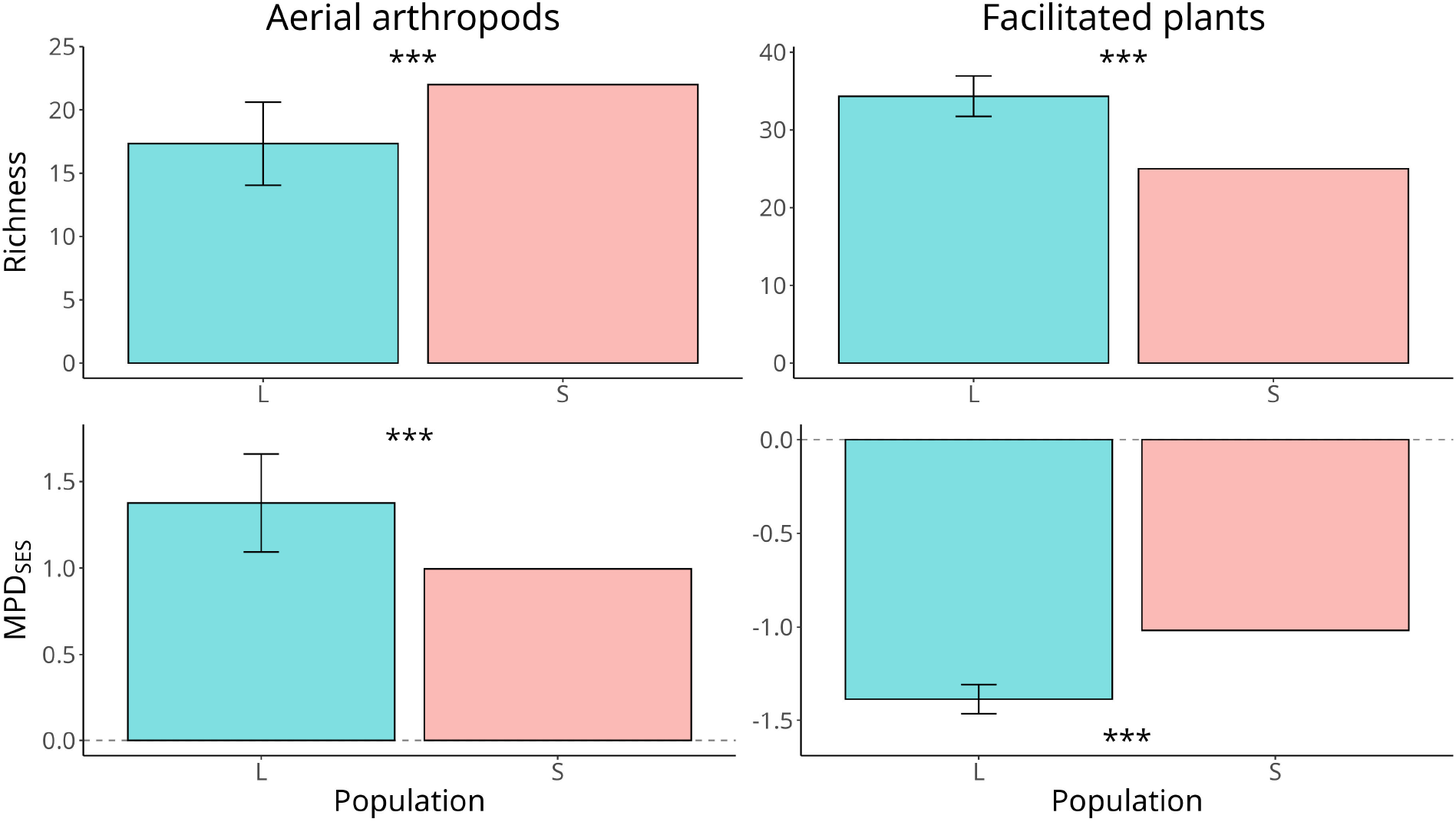
Richness (above) and evolutionary diversity (MPD_SES_; below) of aerial arthropods (left), and facilitated plants (right) in a small population (S, red) of Krascheninnikovia ceratoides and 3 patches of similar size within a large population (L; blue; standard deviation of the 3 patches L1, L2, L3 is shown). Whiskers indicate 95% confidence intervals around the mean. Asterisks indicate significant differences between large and small populations.

## DISCUSSION

Our study aimed to address the extent to which the rarity of plants in a gypsum-rich steppe may contribute or constrain their role in maintaining local diversity through biological interactions, after defining plant rarity as abundance at three different levels: geographic extent, ecological specialization and population size. This integrative perspective reveals that, overall, rare plants tend to support communities of plants and arthropods of similar richness and diversity than more abundant co-occurring plants, while making an important contribution to the uniqueness of such associated communities.

### The effect of host rarity

The richness of the interacting arthropod community was not found to be associated with the geographic extent of plants. While it is well established that biological interactions can influence plant distribution or abundance, especially in extreme environments (Escudero et al. 2015), there appears to be no reciprocity in the sense that the ability to thrive in diverse environments does not translate into higher participation in local interactions or association to more generalist interactuants. Kempel et al. (2020) also provided empirical evidence that the effectiveness of certain interactions, such as competition or response to herbivory, was not always higher in widely distributed compared to rare plants, and concluded that it is essential to consider the ecological context in which the traits associated with interactions evolved in each species. Our results also showed no relationship between the ecological specialisation of plants and the richness and diversity of interactions, as gypsophiles did not stand out either as hosts of a greater diversity or as a distinctive community of arthropods. Santamaría et al. (2018) studied pollinator networks in communities on gypsum soils in the Iberian Peninsula and found a trend towards increased pollinator visits in gypsophiles. It is important to note that our study encompassed all types of interactions, not only pollinators. In the broader context of our study, results suggest that being trapped in edaphic specialisation has no effect on interactions with arthropods.

Most likely, other factors such as plant functional traits, including overall size, leaf shape and floral reward, play a more important role in increasing the number of interactions. Large plants not only increase the diversification of niches or resources offered to aerial visitors, but also a key facilitation mechanism in hot areas: microclimate amelioration (see Fig. 1). Of the 32 plants analysed, trees (e.g. *Pinus halepensis*) and shrubs (e.g. *Artemisia herba-alba, Salvia rosmarinus*) are the ones that accumulated the greatest diversity of aerial visitors (Table 1), and a single tree species distributed throughout the western Mediterranean, *Juniperus thurifera*, was associated with 59% of all the arthropods recorded in the community. As structurally complex plants, trees and shrubs introduce strong environmental heterogeneity into drylands, and provide favourable habitat for small plants and arthropods. Braun et al. (2021) also found that vegetation has direct and indirect physical effects on arthropod-vegetation association patterns in arid ecosystems. This effect was proposed long time ago on the basis of the size and the resource diversity hypotheses (Lawton 1983), and showcases the significant role of larger plants in attracting and facilitating other life forms, regardless of their range or ecological specificity (Alcántara et al. 2024).

With regard to the effect of population size, our study shows contrasted patterns for aerial arthropods and facilitated plants, highlighting the difficulty of generalizing its effect. It is well known that species that are geographically rare or ecologically specialised can also be abundant or even dominant at regional or local scales (Lesica et al. 2006), and this is the case for *K. ceratoides*. Since the two populations sampled showed similar density, this effect is ruled out. In terms of plant-plant interactions, we found that the small population facilitated a less diverse community of plants than all the three patched embebed in the large one, whereas the opposite pattern was found for aerial arthropods. Santamaría et al. (2018) found that increasing patch area led to increased visitation and pollinator richness in fragments of plant communities on gypsum in the central Iberian Peninsula. Similarly, Aizen et al. (2012) showed that patch size and local plant richness positively influenced the number of interactions recorded in isolated hilltop patches in Argentina. Our L patches are not larger than their corresponding S patch, but benefit from the larger extension of the population where they are located. Interestingly, *K ceratoides* is anemophilous and wind dispersed, and for this reason the arthropod community recorded in this study is mainly made up of phytophagous and predators, rather than pollinators. Experimental work with a food web of specialists (parasitoids-aphids) demonstrated that it is not only the host plant area that is relevant for determining the size of food webs, but also the richness of the plant community, as its reduction leads to a drastic simplification of the web due to multitrophic interactions (Petermann et al. 2010). Our results thus suggest that the positive or negative effect of population size may depend on the type of interaction, as found by Andersson et al. (2016) in a set of patches in temperate forests: small patches suffered more pollen limitation, but also escaped herbivore attacks more often.

Rare species have found to make a significant contribution to the functional diversity of the communities or ecosystems in which they occur (Lyons et al. 2005; Mouillot et al. 2013; Leitão et al. 2016; Basile 2022). The role of rare species in ecosystem functioning may be indirect, for example by facilitating the presence of other common and structural species, or through unique functions. It may not even depend on abundance, but on their mere presence (Dee et al. 2019). Simmons et al. (2020) investigated the response of the whole community and its dynamics to the loss of rare individual interactions in real mutualistic interaction networks, and demonstrated that the most vulnerable interactions are the ones that contributed the most to stability in the face of perturbations such as fragmentation. Some of the rarest plant species of this study have been recorded to interact with a limited number of arthropods, and these interactions have been observed to be highly specialised. For instance, the endemic *Vella aspera* was associated with eight arthropods, and they are highly specialised on it, *i*.*e*. these represent ‘rare’ interactions in the community. *K. ceratoides* is another example, as it supported a very small (nine taxa) but unique entomofauna such as the phytophagous *Eurotica distincta* (Homoptera, Psyllidae), which is restricted to this plant and has the same disjunt distribution from the Caucasus to Mongolia (Ribera and Blasco-Zumeta 1998). Interactions of this nature are vital to one of the participants and do not follow the classical nesting pattern of mutualistic networks, where specialist species typically interact more frequently with generalist species (Bascompte et al. 2003).

### Accounting for biological interactions of rare species in conservation policy

Our study combined two elements that have rarely been studied together. On the one hand, while rare organisms are often the focus of attention in conservation biology because they are assumed to be at higher risk (Harnik et al. 2002; Pimm et al. 2014; Harrison and Noss 2017), they are often ignored in studies of ecological processes and functions because they are assumed to have little impact on them. On the other hand, biological interactions are often ignored in conservation biology even though they play a major role in fields like ecological restoration (but see Tylianakis et al. 2010; Valiente-Banuet et al. 2015). If global change push rare hosts to disappear there will be strong reorganisations of the interaction network (rewiring), which could result in losses in the case of the most specialised interactants (Rezende et al. 2007; Bascompte et al. 2019). However, it is difficult to predict the ultimate consequences due to high uncertainties in the responses of the species themselves and the impacts on networks (Tylianakis et al. 2010).

A multispecies experiment has shown that rare plants are less able to cope with climate change than more widely distributed or abundant plants because of their smaller thermal niche breadth (Vincent et al. 2020). However, the analysis of the dynamics of many rare (endemic and specialists) plants of poor substrates such as cliffs has also led to the proposal that they may be more independent of the effects of climate change because of their extremely stable dynamics (Múgica et al. 2024). Land use change is the main driver of changes in biodiversity (IPBES 2019), and the diversity in regions with Mediterranean climate is considered among the most sensitive ones to land use and climate (Newbold et al. 2020). The community studied here is representative of the a Mediterranean steppe biocenosis, which is often fragmented and relegated to patches within a matrix of intensive agricultural use. This threat is exacerbated by climate change, as water availability is one of the major constraints on the development of the plants that inhabit it. Two of the species with the highest arthropod specialisation are protected in different catalogues due to their global or regional rarity: *V. aspera* (Wild species under special protection in the region of Aragon, included in Annexes II and IV of the European Habitats Directive) and *K. ceratoides*: Vulnerable at regional level). In this context of double vulnerability (habitat and host plants), the loss of plant species is likely to be highly detrimental due to the high frequency of very specific pairwise associations with arthropods, and their relict nature (Ribera & Blasco-Zumeta 1998).

Although our results should be considered a case study because they are restricted to a steppe community, gypsum soils are distributed on all continents and host numerous rare edaphic endemisms (Ortiz-Brunel et al. 2023). Our integrative approach to the multifaceted concept of rarity did not provide evidence for a pattern between host rarity and the abundance and diversity of interacting organisms. What they did show is the enormous diversity and sometimes high specificity of interactions that some rare plants undergo, highlighting their probably underestimated importance for the functioning of the ecosystems in which they occur.

## Notes

https://doi.org/10.20350/digitalCSIC/15101

### Competing Interest Statement

The authors have declared no competing interest.

### Funding

- Fundación BBVA: Perdiendo la red. El papel de plantas amenazadas en la conservación de la diversidad global. PERDIVER
- CYTED: Interacciones ecológicas como indicadoras de gestión en áreas naturales protegidas (INTERGAP)

